## Supplemental Data for "Chronic RNA G-quadruplex Accumulation in Aging and Alzheimer’s Disease"

**Supplementary Data Table 1.** Antibodies used in the study and their respective dilutions and sources.

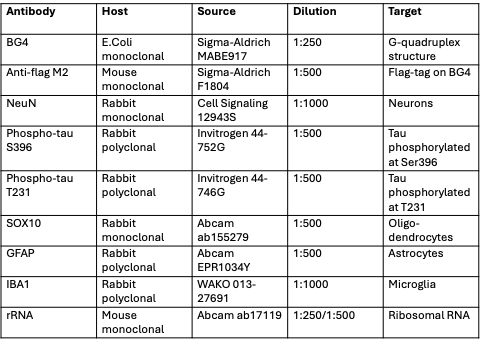

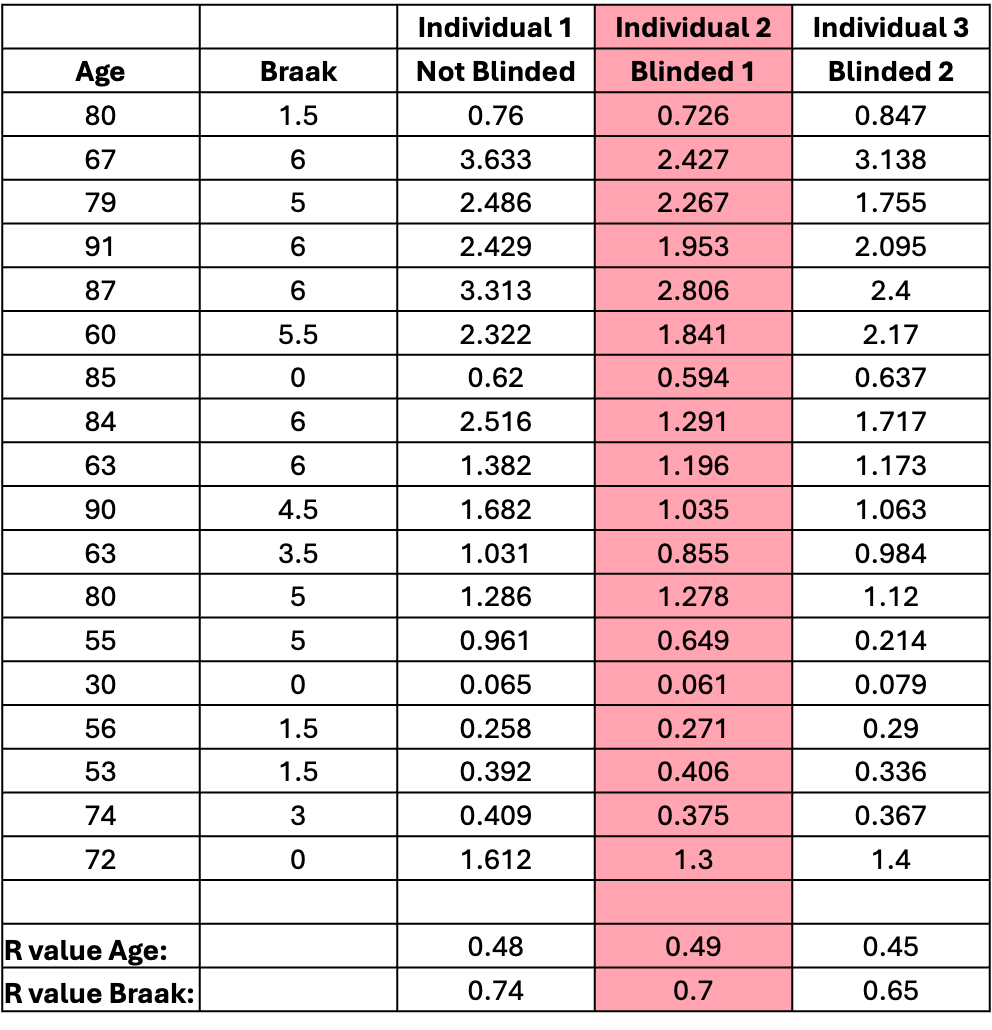
**Supplementary Data Table 2.** Quantification of BG4 stain by % Area in the OML performed by 3 different individuals, 2 of which were completely blind to the case information. Highlighted column is the quantification included in the main text. All 3 R values are highly significant.

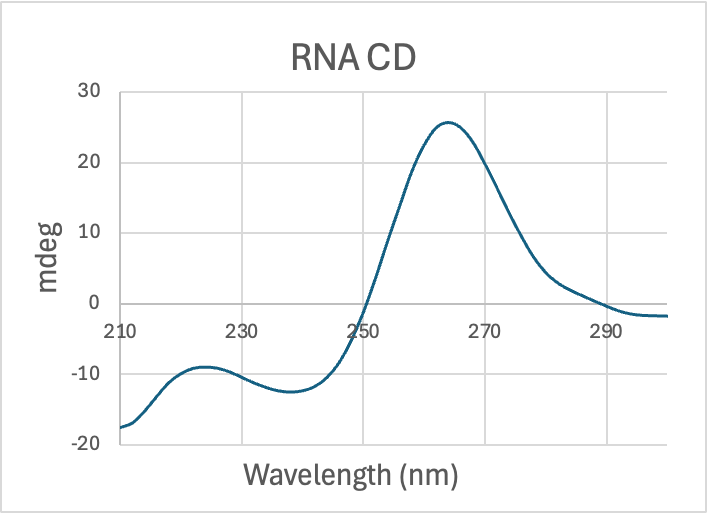

**Supplementary Data Fig 1.** Circular dichroism spectra showing G-quadruplex structure of RNA sequence able to most efficiently template tau fibrillation in Zwierzchowski-Zarate et al (see Introduction of main text for description) (5'-CGGGCGGCGGGGGGGCCCGGGCGGCGGGGGGGCCCGGGCG-3').

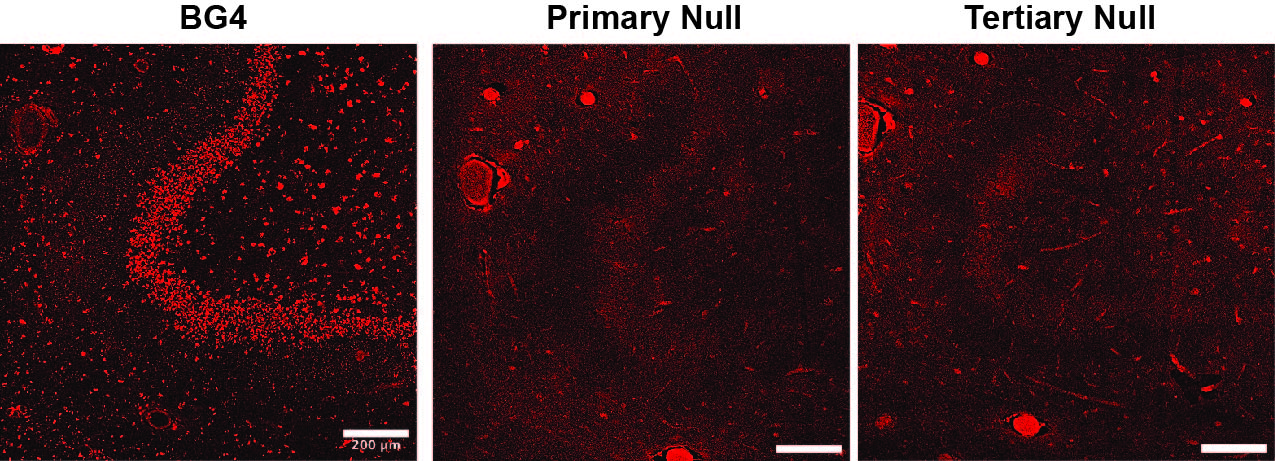

**Supplementary Data Fig 2. Antibody null controls.** (left) Complete BG4 staining for comparison. (center) BG4 primary null in incubated with intermediate and tertiary antibodies only. (right) BG4 tertiary null is incubated with primary and intermediate antibodies only.

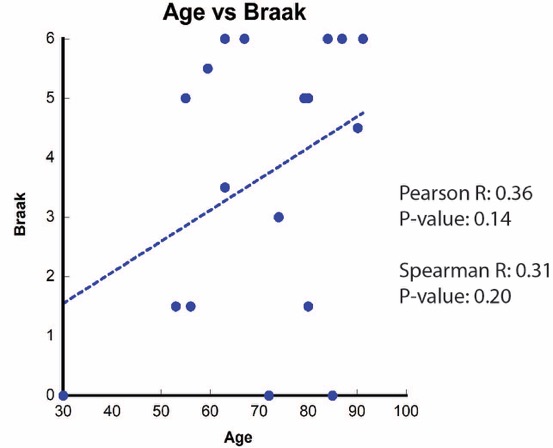
.

**Supplementary Data Fig 3.** Very weak correlation was observed between age and Braak stage.

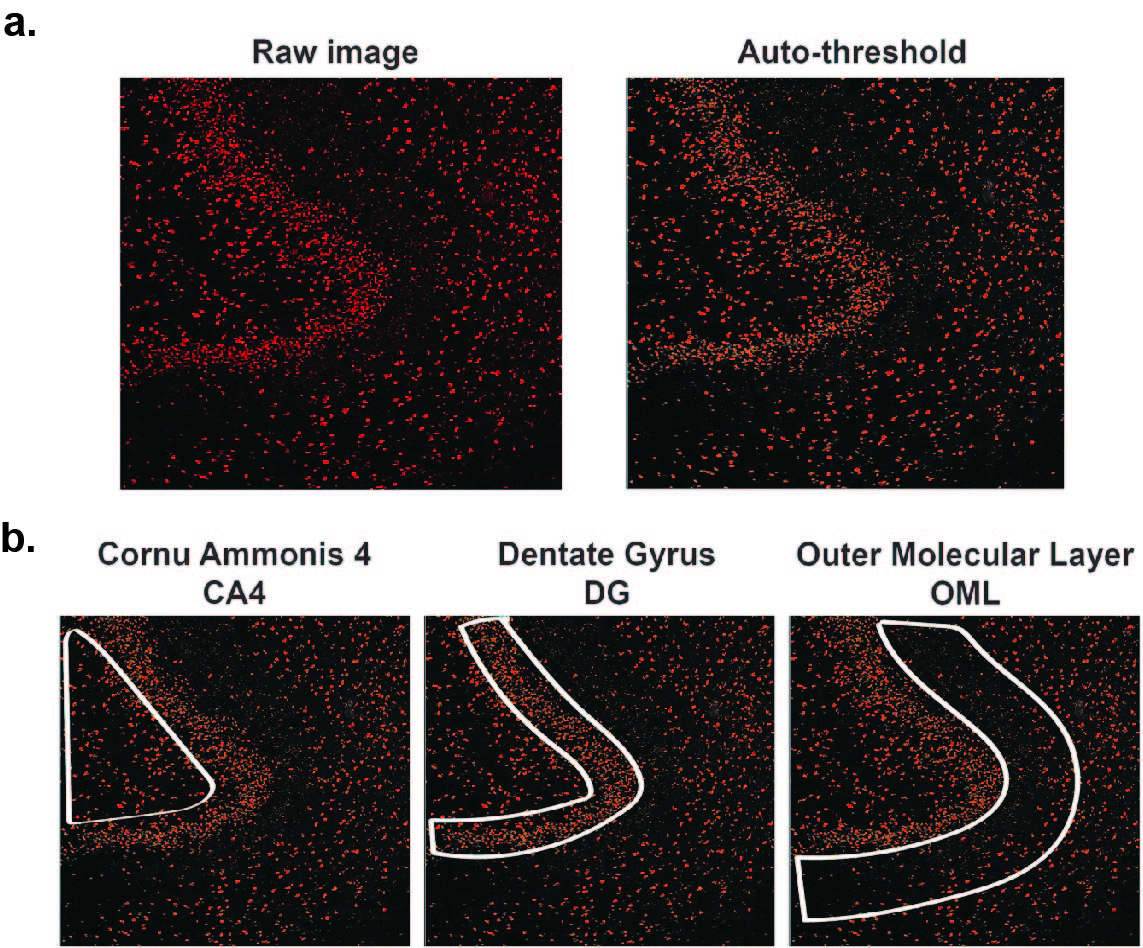

**Supplementary Data Fig 4. ImageJ quantification. a,** Unedited confocal image on left. Auto-threshold set by the program ImageJ applied in the BG4 channel on right. **b,** Representative regions of interest used in quantification for each region of the hippocampus.

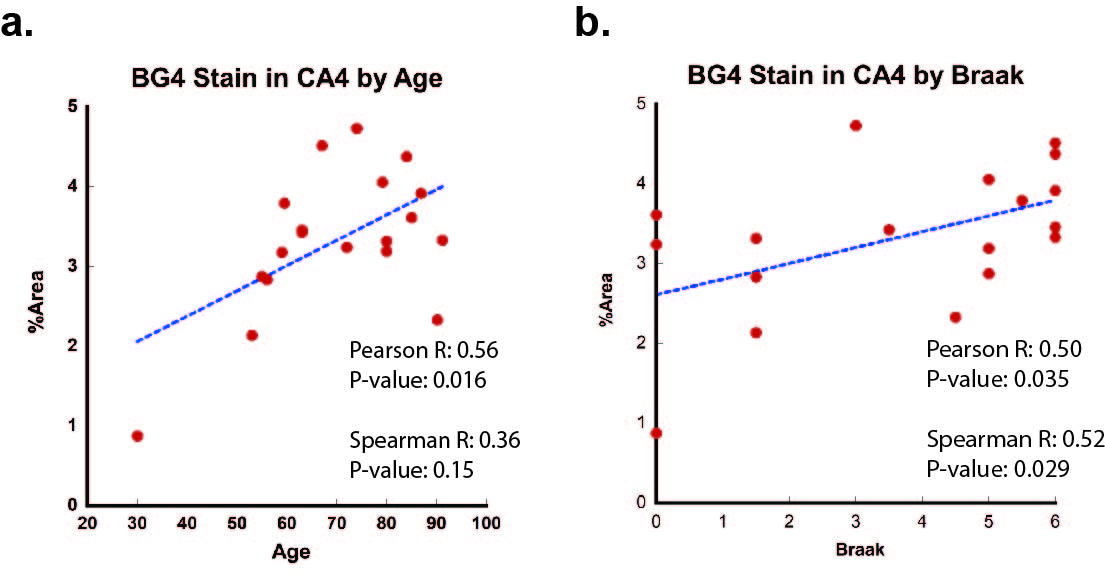

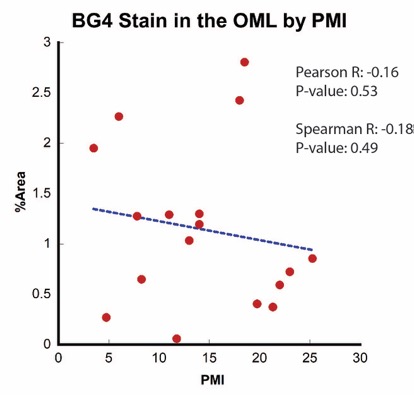
**Supplementary Data Fig 5. a,** BG4 stain in the CA4 region exhibits positive Pearson correlation with age, but no Spearman correlation. **b,** BG4 exhibits significant positive correlation with Braak stage in CA4 region.

**Supplementary Data Fig. 6.** No significant correlation observed between BG4 stain and PMI

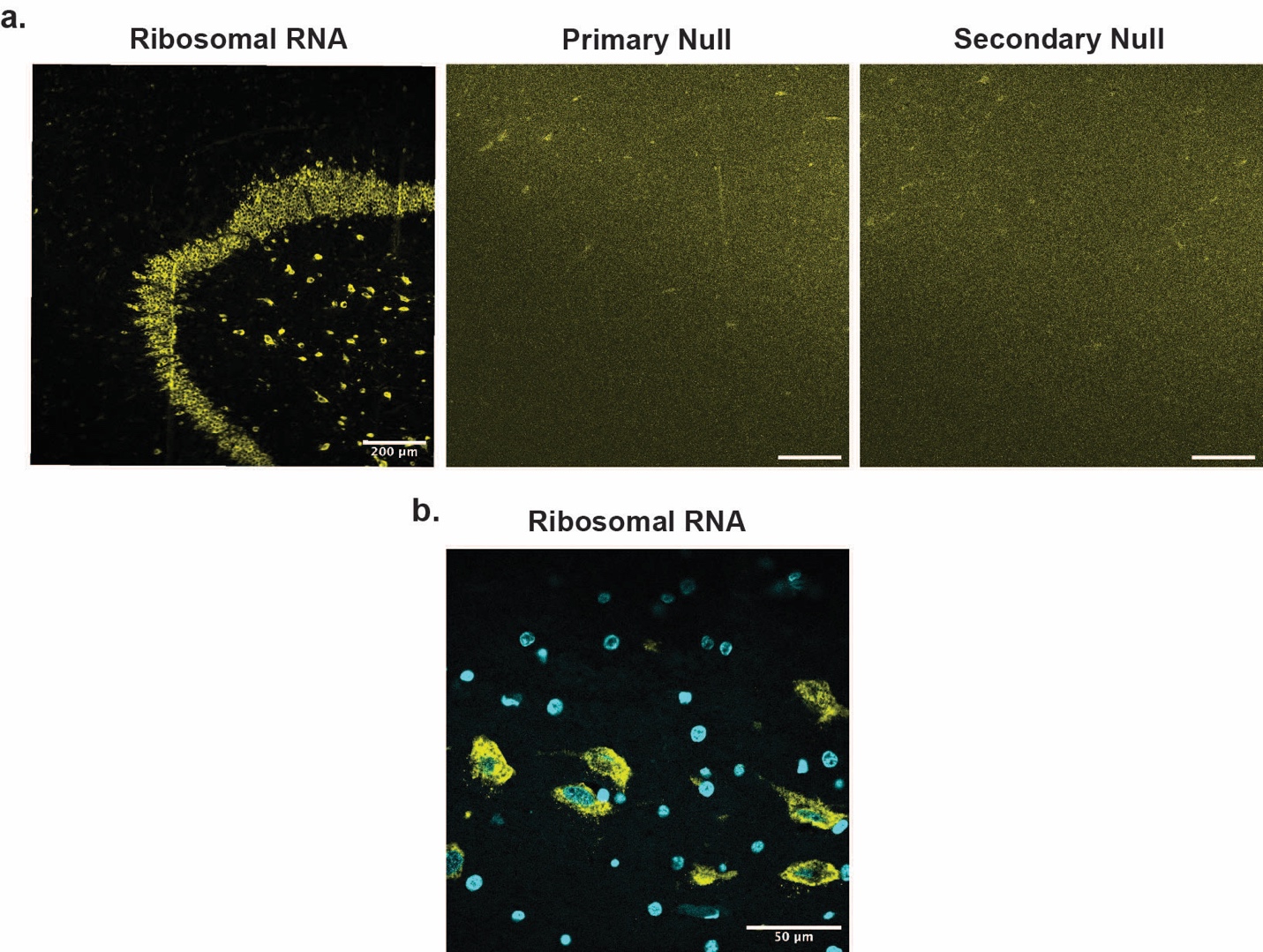

**Supplementary Data Fig 7. Ribosomal RNA antibody. a,** (left) Complete rRNA staining in hippocampal region of a young control individual, using the same staining protocol as for BG4. (center) rRNA antibody primary null incubated with secondary only. (right) rRNA antibody secondary null incubated with primary only. **b,** 60X magnification of rRNA antibody.
